# A distributed pallial circuit links sensory and bodily representations to aversive motivational value in the goldfish dorsomedial pallium

**DOI:** 10.64898/2026.08.04.742847

**Authors:** Carmen Salas-Peña, Blanca Quintero, Adrián Chinarro, Antonia Gómez, Daniel Lozano, Jesús M. López, Fernando Rodríguez, Nerea Moreno, Cosme Salas

## Abstract

Understanding how neural circuits transform sensory and bodily signals into motivational states and adaptive behavior is a central problem in neuroscience. In teleost fish, the dorsomedial telencephalon (Dm) is a key pallial region implicated in both sensory processing and aversive behavior, yet whether these functions arise from a functionally uniform region or from interactions among specialized pallial domains has remained unknown. Here we show that the teleost dorsomedial telencephalon exhibits a previously unrecognized functional organization in which distinct but interconnected pallial domains perform complementary computations that progressively transform multimodal sensory and bodily representations into aversive motivational value and adaptive behavioral control. Wide-field voltage-sensitive dye imaging revealed that tactile, auditory, and gustatory stimuli evoke spatially organized, modality-specific activity exclusively within the caudal subdivision of Dm (Dmc), whereas the rostral subdivision (Dmr) showed little or no sensory responsiveness. In contrast, focal intracerebral microstimulation demonstrated that activation of Dmr, but not Dmc, is sufficient to generate robust, flexible, and reversible conditioned place avoidance, identifying Dmr as a pallial node causally involved in the assignment of negative motivational value. Anatomical tracing revealed a circuit in which sensory and bodily-related inputs converge onto Dmc, are relayed intrapallially to Dmr, where they are transformed into an aversive motivational signal before being conveyed to hypothalamic and brainstem centers involved in autonomic and behavioral regulation. Immunohistochemical analyses confirmed the pallial identity of both subdivisions and their distinct rostrocaudal organization, while providing no evidence that Dm corresponds to a classical pallial amygdaloid territory. This functional architecture more closely resembles the distributed organization of mammalian corticolimbic networks than either a unitary pallial amygdala or a neocortical sensory hierarchy, suggesting that the transformation of sensory and bodily representations into motivational control may represent a conserved organizational feature of the pallium that emerged early during vertebrate evolution.

**Short abstract / Significance statement:** This study shows that the teleost dorsomedial pallium is organized into complementary functional domains that dissociate multimodal sensory representation from negative motivational processing while forming an interconnected pallial circuit associated with adaptive behavioral control. Our findings reveal a distributed pallial organization resembling mammalian corticolimbic architectures and provide a new framework for understanding the evolution of vertebrate pallial function.

## INTRODUCTION

Understanding how sensory and interoceptive information is transformed into motivational states, emotional responses, and adaptive behavior is a central problem in neuroscience (Damasio, 1999; Mesulam, 1998; Pessoa, 2013; Cisek, 2019). These transformations rely on forebrain circuits that integrate sensory inputs with bodily signals and assign motivational value to biologically relevant events, thereby guiding adaptive behavior (Mogenson, Jones & Yim, 1980; Swanson, 2000; Pessoa, 2017). While these processes have been extensively investigated in mammals, their neural organization in other vertebrate groups remains less understood. Comparative studies across vertebrates are therefore essential for identifying the organizational principles through which sensory and bodily information are transformed into motivational value, emotional states and adaptive behavior.

In teleost fish, the telencephalic pallium plays a major role in emotional and motivational behavior (Rodríguez et al., 2002; Salas et al., 2003; Portavella et al., 2004; Overmier & Hollis, 1990; O’Connell & Hofmann, 2011). Within the pallium, the medial division (Dm) has been consistently implicated in aversively motivated learning and defensive responses. Lesion studies in goldfish and other teleost species have shown that damage to Dm disrupts avoidance learning and conditioned autonomic responses associated with negative outcomes (Portavella et al., 2004; Broglio et al., 2005; Martín et al., 2011; Lal et al., 2018). These findings identify Dm as a key telencephalic component of the neural circuitry underlying aversive motivational processing and aversive behavior in teleost fish. At the same time, electrophysiological and imaging studies have reported sensory-evoked activity within restricted portions of Dm (Ocaña et al., 2026; Prechtl et al., 1998; Trinh et al., 2026), suggesting that this pallial region may also participate in sensory processing. These apparently disparate sensory and motivational functions have remained difficult to reconcile and have contributed to competing interpretations of the functional and evolutionary significance of Dm. Together, these observations raise the question of how the diverse sensory and motivational functions attributed to Dm are functionally organized within this pallial region.

Several anatomical studies have proposed that Dm is not a homogeneous structure but can be subdivided along its rostrocaudal axis into caudal and rostral domains based on cytoarchitectural features, histochemical markers, and patterns of connectivity (Castro et al., 2003; Northcutt, 2006; Yamamoto et al., 2007; Yáñez et al., 2021; Porter and Mueller, 2022). In goldfish, this subdivision can be identified reliably using macroscopic landmarks together with cytoarchitectural and histochemical criteria (Northcutt, 2006). These anatomical observations suggest that Dm may be composed of distinct functional domains rather than a functionally uniform pallial region. However, the functional significance of this rostrocaudal organization of Dm remains largely unexplored, particularly regarding whether these subdivisions make distinct contributions to sensory processing, negative motivational value assignment, and the integration of these functions within pallial circuits.

Here we investigated whether the caudal (Dmc) and rostral (Dmr) subdivisions of the goldfish Dm support distinct functional roles in sensory processing and aversive motivational control. To address this question, we combined wide-field optical imaging of sensory-evoked activity, intracerebral microstimulation-induced place aversion, histochemical analyses, and anatomical tracing. This multimodal approach allowed us to compare sensory responsiveness, behavioral effects of local activation, and circuit organization in Dmc and Dmr, and to determine how these two pallial domains contribute to the functional organization of sensory processing and aversive valuation, and how these complementary functions are integrated through pallial circuitry.

## RESULTS

### Anatomical and neurochemical parcellation of the dorsomedial pallium identifies two rostrocaudally distinct domains

The dorsomedial pallium (Dm) of the goldfish comprises two rostrocaudally distinct domains, a caudal (Dmc) and a rostral (Dmr) (Northcutt, 2006). These two domains can be reliably identified based on macroscopic landmarks and cytoarchitectural features (Figure 1). From the dorsal surface of the telencephalon, Dmc and Dmr appear as adjacent pallial protrusions separated by shallow valleculae, allowing their boundaries to be consistently delineated across animals (Figure 1A). In addition to these macroscopic features, Dmc and Dmr differ in overall cytoarchitecture and neurochemistry, consistent with previous anatomical and hodological studies in teleost (Northcutt, 2006; Castro et al., 2003, 2006; Yáñez et al., 2021; Ganz et al., 2014).

**Figure 1.**
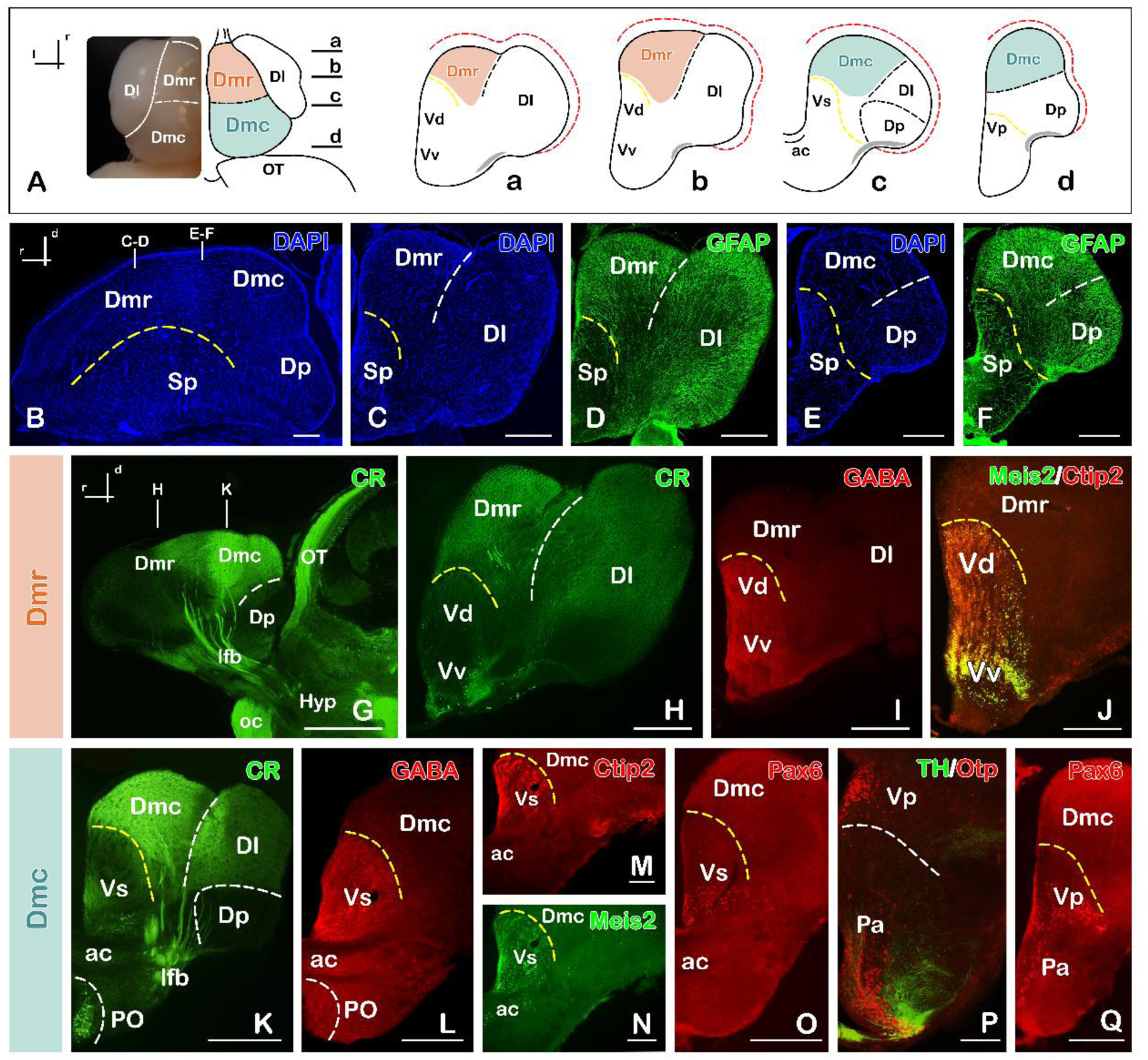
Neuroanatomical boundaries and regionalization of the goldfish dorsomedial pallium along the rostrocaudal axis. **A**. Dorsal view of the goldfish telencephalic hemispheres showing a photomicrograph (left) and the corresponding schematic drawing (right). The transverse sections (a–d), taken at the rostrocaudal levels indicated in the dorsal scheme, illustrate the subdivision of the dorsomedial pallium (Dm) into a rostral domain (Dmr, orange) and a caudal domain (Dmc, blue). Sagittal (**B**) and transverse (**C–F**) sections stained with DAPI and glial fibrillary acidic protein (GFAP). The rostrocaudal levels of the transverse sections (**C–D** and **E–F**) are indicated in the sagittal section shown in B. Sagittal (**G**) and transverse (**H–Q**) sections illustrating the distribution of calretinin (CR), GABA, Meis2, Ctip2, Pax6, tyrosine hydroxylase (TH), and Otp immunoreactivity. The color code for each marker is shown in the upper corner of the corresponding panel. In the sagittal sections (**B, G**), the rostrocaudal and dorsoventral axes are indicated in the upper left corner. Scale bars, 100 μm. The dashed red line in every scheme represents the tela choroidea. The dashed yellow line in every image and scheme represents the pallial-subpallial boundary. *Abbreviations*: ac, anterior commissure; Dl, lateral part of the dorsal telencephalic area; Dlv, ventral division of the lateral part of the dorsal telencephalic area; Dm, medial part of the dorsal telencephalic area; Dmc, caudal division of the medial part of the dorsal telencephalic territory; Dmr, rostral division of the medial part of the dorsal telencephalic territory; Dp, posterior part of the dorsal telencephalic territory; Hyp, hypothalamus; lot, lateral olfactory tract; lfb, lateral forebrain bundle; oc, optic chiasma; OT, optic tectum; Pa, paraventricular hypothalamus; PO, preoptic area; Sp, subpallium; Vd, dorsal part of the ventral telencephalic area; Vp, postcommissural part of the ventral telencephalic area; Vv, ventral part of the ventral telencephalic area; Vs, supracommissural part of the ventral telencephalic area.

DAPI nuclear staining revealed clear differences in the cellular organization and packing between Dmc and Dmr (Figure 1B). Along the rostrocaudal axis, the cells in the external portion of Dm were predominantly arranged parallel to the pial surface, although cell packing was denser rostrally than caudally. Comparative cytoarchitectonic analysis using GFAP immunostaining to visualize the radial glial processes defining this domain revealed a characteristic wedge-shaped cellular organization extending from the pial surface toward the interior. At rostral levels, Dmr was restricted to a compact dorsomedial territory composed of densely packed cells extending radially from the ventricular zone. At caudal levels, this territory expanded dorsally while maintaining its close association with GFAP-positive radial glial processes (Figure 1C–F).

To further characterize Dm, we analyzed the expression of conserved neuronal markers previously used to define pallial and subpallial territories in teleost (Castro et al., 2006; Tibi et al., 2023; Fig. 1G–Q). Immunostaining for the calcium-binding protein calretinin (CR) was particularly useful for identifying Dm and its internal regionalization. In agreement with previous studies (Northcutt, 2006; Castro et al., 2006; Porter & Mueller, 2020; Folgueira & Clarke, 2024), CR immunoreactivity revealed pronounced rostrocaudal differences within Dm (Figure 1G). A dense CR-positive fiber innervation was observed in Dmc, whereas CR labeling was substantially weaker in Dmr (Figure 1H,K). This differential pattern also helped delineate the boundaries between Dm and adjacent pallial and subpallial regions, where CR innervation was sparse or absent.

The pallial identity of Dm was further supported by markers that distinguish pallial from subpallial territories (Tibi et al., 2023; Ganz et al., 2012, 2014). GABA immunostaining was absent throughout Dm, clearly delineating the pallial-subpallial boundary (Fig. 1I,L). This boundary was further supported by the expression of the subpallial markers Meis2 and Ctip2, which were restricted to ventral telencephalic territories and did not extend into either Dmr (Figure 1J) or Dmc (Figure 1M,N).

Finally, we examined the expression of conserved molecular markers associated with amygdaloid territories (Biechl et al., 2017; Armbruster et al., 2025; Porter & Mueller, 2020), including Otp, Pax6, and tyrosine hydroxylase (TH). These markers were selectively expressed in the supracommissural (Vs) and postcommissural (Vp) nuclei of the ventral area, corresponding to the extended amygdala of the goldfish (Figure 1O–Q). In contrast, the adjacent Dmc lacked this molecular profile, indicating that it does not share the molecular identity of amygdaloid territories.

Together, these anatomical, cytoarchitectural, and molecular features define Dmr and Dmc as two distinct pallial domains. These complementary landmarks enabled their consistent identification across all animals, delineated their boundaries with adjacent pallial territories, and provided the anatomical framework for the subsequent functional and connectivity analyses.

### Sensory-evoked activity is restricted to the caudal subdivision of Dm (Dmc)

To determine how sensory information is organized within Dm, we examined tactile-, auditory-, and gustatory-evoked responses using in vivo wide-field voltage-sensitive dye imaging (Figure 2). Sensory stimulation reliably evoked neural activity within Dm; however, this activity was consistently restricted to the caudal subdivision (Dmc) in all animals examined. In contrast, the rostral subdivision (Dmr) showed little or no sensory responsiveness under the present recording conditions.

**Figure 2.**
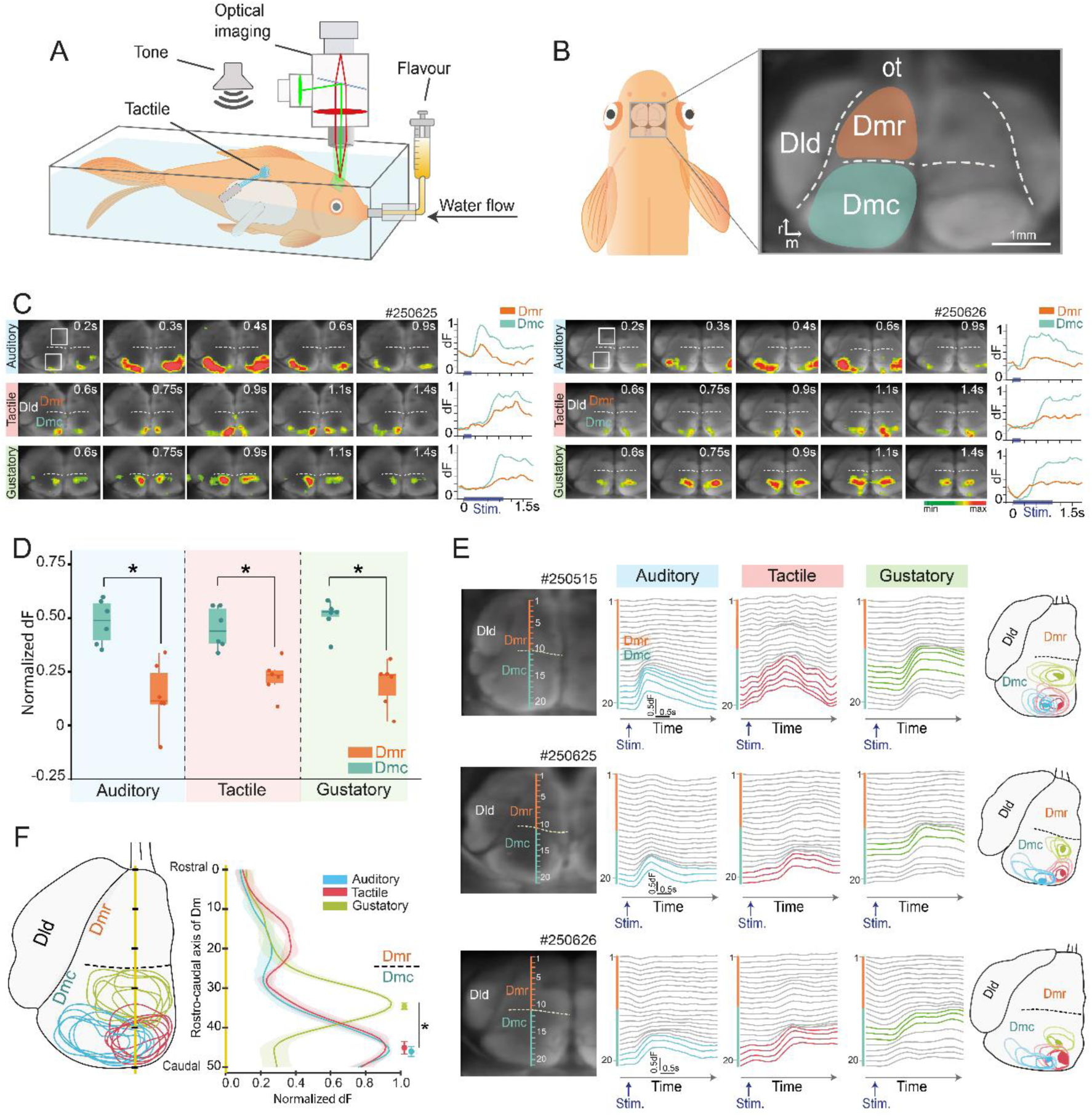
Pallial activity evoked by sensory stimulation in the goldfish Dm. **A.** Experimental setup for wide-field voltage-sensitive dye imaging of sensory-evoked activity. **B**. Schematic representation of the dorsal telencephalon of the goldfish and optical imaging field. The right panel shows a representative fluorescence image of the dorsal telencephalic surface after staining with the voltage-sensitive dye, highlighting the relative position of the rostral (Dmr) and caudal (Dmc) subdivisions of Dm and adjacent pallial regions (Dld). **C.** Spatiotemporal patterns of pallial activation evoked by auditory, tactile, and gustatory stimulation. Sequential pseudocolor frames from in vivo optical imaging in two representative animals show the time course of sensory-evoked fluorescence changes (dF) in the pallium after stimulus onset, with fluorescence changes at each pixel color-coded according to the scale bar. Time elapsed from stimulus onset is indicated in each frame. Traces on the right show the time course of the normalized VSD signal extracted from a 15×15 pixels ROIs (white squares) centered in Dmc and Dmr for the representative animal shown in each row. Across all sensory modalities, evoked activity was largely confined to Dmc, whereas Dmr exhibited little or no response. Note that sensory-evoked activity was consistently restricted to Dm and did not extend into the adjacent pallial subdivision Dld. **D.** Sensory-evoked activity in Dmc and Dmr across sensory modalities. Mean normalized evoked activity (dF) was quantified within each ROI during the 1.680 ms (280 frames) following stimulus onset for auditory, tactile, and gustatory stimulation. Responses were significantly greater (asterisks) in Dmc than in Dmr across all sensory modalities. Each dot represents an individual animal (n = 6). **E.** Spatiotemporal propagation of sensory-evoked activity within Dm in three representative animals. Right, sequential (superimposed) outlines illustrating the spatial propagation of sensory-evoked activity from response onset to peak activation for auditory (blue), tactile (red), gustatory (green) stimulation on a drawing of the left telencephalic hemisphere of each subject. For each sensory modality, contours depict the spatial extent of the active area at five successive time points: the earliest detectable response (25% of peak activity), the peak response, and three intermediate time points. Contour shading progresses from dark to light, representing the temporal evolution from the early response to peak activation. Filled contours indicate the earliest detectable responses, revealing modality-specific sites of activation onset within Dmc. Left, activity profiles extracted from 21 consecutive sampling regions spanning the rostrocaudal axis of Dm are shown over the course of the sensory-evoked response for auditory, tactile and gustatory stimulation. Color-coded traces correspond to sampling regions whose activity exceeded 75% of the normalized peak response. Activity profiles indicate that suprathreshold responses are consistently confined to the caudal portion of Dm for all modalities. **F**. Summary of the topographic organization of sensory representations within Dm. Left, superimposed contours of peak-evoked activity illustrating the spatial arrangement of the auditory (red), tactile (blue), and gustatory (green) representations within Dm. Data from all fish (n = 6) were spatially aligned and overlaid in a schematic representation of the left telencephalon using the external borders and the ypsiloniformis sulcus as landmarks. Right, rostrocaudal activity profiles for each modality were generated by averaging across animals the normalized activity values extracted at the time of the peak response from a line profile spanning the rostrocaudal axis of Dm (50 consecutive sampling points). Symbols above the curves indicate the mean (± SEM) location of maximal activity for each modality along the rostrocaudal axis of Dm, revealing a more rostral peak response to the gustatory stimulus than to the auditory or tactile stimulus. Asterisk denotes statistically significant differences.

Analysis of the spatiotemporal activity maps revealed a marked spatial segregation of sensory-evoked activity between the Dmc and Dmr domains (Figure 2C). To quantify this pattern, we compared the mean fluorescence signal within two 15×15 pixels ROIs centred on the Dmc and Dmr regions over a 1.680 ms period following stimulus onset. Quantitative analysis confirmed that mean evoked activity was significantly greater in Dmc than in Dmr across all three sensory modalities (n = 6; paired-sample t tests: tactile, t(5) = 5.961, p = 0.002; auditory, t(5) = 6.823, p = 0.001; gustatory, t(5) = 13.702, p < 0.001; Figure 2D). These results indicate that tactile-, auditory-, and gustatory-evoked responses were consistently confined to Dmc and spread throughout this subdivision, whereas little or no propagation into Dmr was observed.

### Topographic and modality-specific organization of sensory representations within Dmc

Having shown that sensory responses are confined to Dmc, we next examined the spatial organization of sensory-evoked activity within this pallial subdivision. Sensory-evoked activity formed reproducible spatial activation patterns in which tactile, auditory, and gustatory stimuli activated distinct regions within Dmc (Figure 2E,F).

To characterize the spatial and temporal organization of these activity patterns, we analyzed the spatiotemporal dynamics of sensory-evoked activity following stimulus onset by tracking the propagation of the active area from the early to the peak response for each sensory modality. The earliest sensory-evoked responses revealed a clear spatial organization within Dmc (see filled contours in Figure 2E). Gustatory responses first were detected in a rostral sector of Dmc, whereas tactile and auditory responses first appeared in its caudal region. Within this caudal domain, tactile responses were located more medially, whereas auditory responses occupied a more lateral position. As activity progressed toward its peak, tactile-evoked activity progressively expanded and developed partial spatial overlap with the auditory representation. In contrast, gustatory-evoked activity partially overlapped medially with tactile-evoked activity but remained spatially distinct from the auditory representation.

To characterize more precisely the spatial organization of these sensory-evoked activity patterns, we analyzed the distribution of sensory-evoked activity along the rostrocaudal axis of Dm using a series of 21 consecutive positions spanning this axis (Figure 2E). This analysis identified the regions exhibiting the strongest activity and quantified the spatiotemporal propagation of responses for each of the three sensory modalities, revealing distinct spatial and temporal dynamics across modalities. Across all modalities, the strongest activity (≥75% of the peak response, colored traces in Figure 2E) remained confined to the Dmc subdivision, corresponding to the caudal half of the rostrocaudal axis of Dm, throughout the evoked response. However, the spatiotemporal propagation patterns differed among the three sensory modalities. Gustatory-evoked activity consistently occupied a more rostral position, whereas tactile- and auditory-evoked activity was centered in more caudal regions of Dmc (Figure 2E). In addition, onset latencies (defined as the time to reach 25% of the peak amplitude) differed significantly across sensory modalities (repeated-measures ANOVA: F(2,10) = 6.894, p = 0.013). Bonferroni-corrected post hoc comparisons revealed that auditory-evoked activity exhibited shorter onset latencies than gustatory-evoked activity (p = 0.036). A similar trend was observed for the comparison between auditory- and tactile-evoked activity, although this difference did not reach statistical significance (p = 0.065). No significant differences were found between tactile- and gustatory-evoked responses (p = 1). These findings suggest a faster temporal course of auditory responses.

To compare the spatial distribution of evoked activity along the rostrocaudal axis of Dm across sensory modalities, normalized activity values at the time of the peak response were extracted from a line profile spanning the rostrocaudal axis, yielding 50 consecutive sampling points. These values were averaged across subjects within each modality to generate rostrocaudal activity profiles (Figure 2F), which were then used to statistically compare the positions of maximal activity. For all three sensory modalities, the position of maximal evoked activity along the rostrocaudal axis was located within Dmc. However, the rostrocaudal position of the activity peak differed significantly across sensory modalities (repeated-measures ANOVA: F(2,10) = 86.255, p < 0.001). Bonferroni-corrected post hoc comparisons revealed that both tactile- and auditory-evoked activity peaks were located significantly more caudally than the gustatory-evoked activity peak (both p < 0.001), whereas no significant difference was observed between the tactile and auditory modalities (p = 1). This spatial organization was further supported by the superimposed peak response maps from the three sensory modalities, which revealed a consistent topographic arrangement of sensory-evoked activity within Dmc (Figure 2F). Although the three sensory representations partially overlapped, each modality occupied a characteristic preferential territory within Dmc. Notably, the somatosensory representation occupied an intermediate position, partially overlapping both auditory and gustatory representations, whereas the latter exhibited little direct overlap with one another. Together, these imaging data demonstrate that Dmc contains a reproducible topographic organization of modality-specific sensory representations, whereas Dmr lacks detectable sensory responsiveness under the present recording conditions.The central position of the somatosensory representation, bridging the auditory and gustatory territories, suggests that body-related sensory information may occupy a key position within the integrative organization of Dmc.

### Focal microstimulation of Dmr, but not Dmc, induces conditioned place avoidance

Having established that sensory-evoked activity is selectively confined to Dmc, we next asked whether either rostrocaudal subdivision of Dm contributes causally to aversive motivational behavior. To test this hypothesis, we combined chronic focal intracerebral microstimulation of either Dmr or Dmc with a conditioned place avoidance paradigm in freely moving goldfish using a serial A–B–A reversal design, in which the stimulation-paired compartment was alternated across successive phases (Figure 3A).

**Figure 3.**
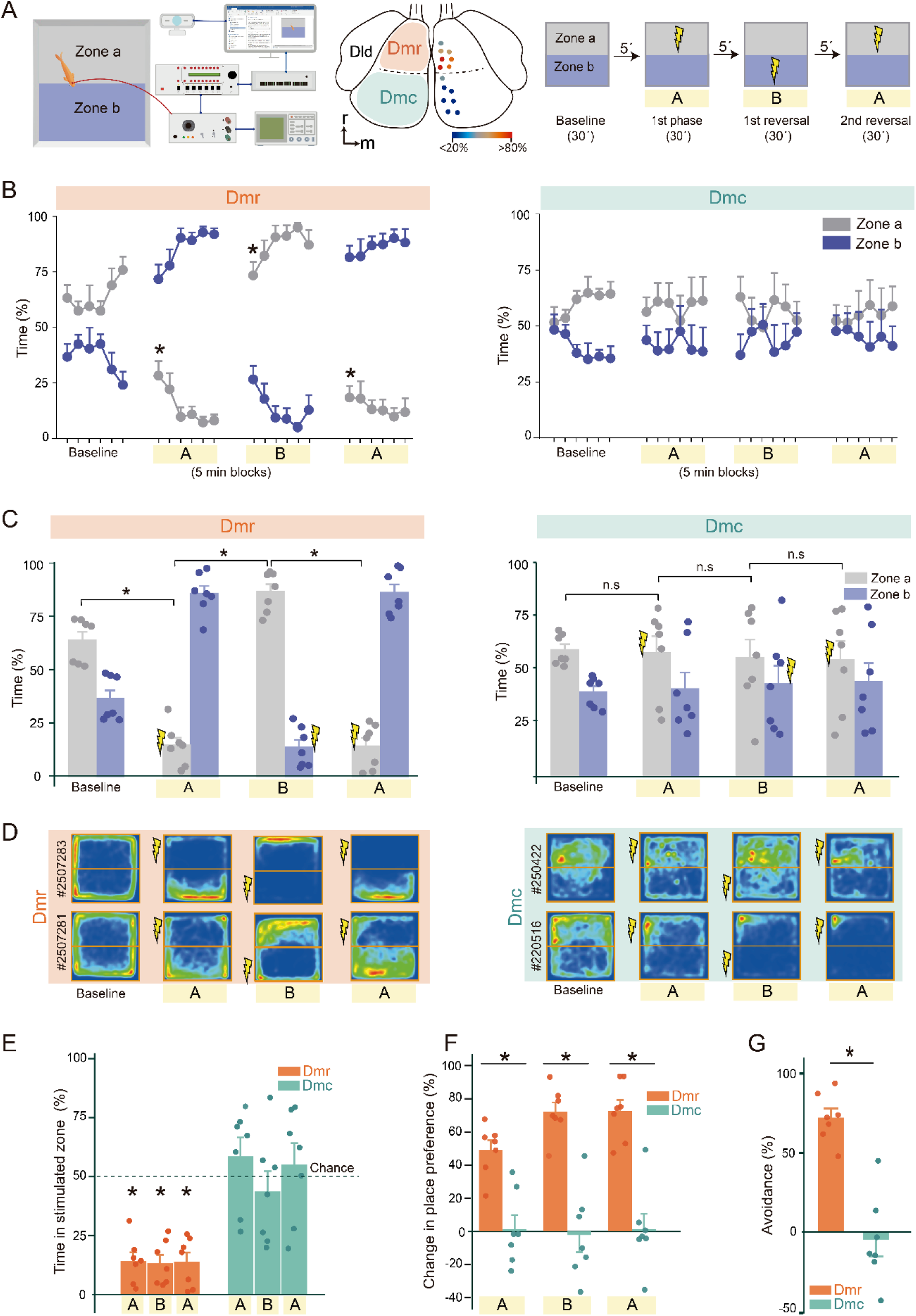
Electrical microstimulation of the Dmr region produces conditioned place aversion. **A.** Schematic representation of the experimental setup and procedure. Left, diagram of the two-compartment arena and recording and stimulation equipment. Center, dorsal view of the goldfish telencephalon showing the location of stimulation electrodes implanted in Dmr and Dmc; color coding indicates the magnitude of avoidance induced by stimulation at each site. Right, timeline of the conditioned place avoidance protocol, including baseline and three successive phases (A–B–A design) comprising two reversals. The yellow symbol marks the compartment where intracerebral microstimulation was delivered. **B.** Mean percentage of time spent in each compartment across baseline and stimulation phases for the group of animals with electrodes implanted in Dmr (left) or Dmc (right). Symbols represent the percentage of time spent in each compartment during consecutive 5-min blocks. For each subject, zone A was defined as the compartment preferred during baseline. During the first stimulation phase, animals received stimulation upon entering the initially preferred compartment; during subsequent phases, the stimulation-paired compartment was reversed. Asterisks denote statistically significant differences in the time spent in the initially preferred compartment between the final 5-min block of one phase and the initial 5-min block of the subsequent phase. **C.** Mean percentage of time spent in each compartment during the full 30-min duration of each phase for Dmr and Dmc groups. Asterisks indicate statistically significant differences. **D.** Occupancy heat maps from two representative animals per group illustrating spatial behavior across baseline and stimulation phases. Animals stimulated in Dmr show clear avoidance of the stimulation-paired compartment, whereas animals stimulated in Dmc do not. **E.** Mean percentage of time spent in the stimulation-paired compartment during each stimulation phase for Dmr and Dmc groups. The dashed line indicates chance level (50%). Asterisks indicate statistically significant differences relative to chance. **F.** Change in place preference between successive phases, showing significantly larger preference shifts in animals with electrodes implanted in Dmr compared with Dmc across all phases. Asterisks denote statistically significant between-group differences. **G.** Mean avoidance index averaged across the three stimulation phases for Dmr and Dmc groups. Asterisk denotes statistically significant between-group differences.

During the baseline phase, animals exhibited a slight spontaneous preference for one compartment of the arena (hereafter referred to as the preferred compartment). Nine of the fourteen tested fish preferred the light-blue compartment, spending an average of 64.5% of their time there, whereas the remaining five preferred the pink compartment, spending an average of 55.98% of their time in that compartment. No significant between-group differences were observed in the percentage of time spent in each animal’s preferred compartment during the baseline period (t(12) = 0.784, p = 0.448; Figure 3B,C). Thus, both groups displayed comparable baseline spatial preferences before the onset of intracerebral stimulation.

Animals receiving focal microstimulation in Dmr rapidly reversed their initial place preference, consistently avoiding the stimulation-paired compartment throughout the conditioning phase (Figure 3B). Reversal of the stimulation contingency produced corresponding reversals of place preference in subsequent phases, demonstrating that the behavioral effect was robust, flexible, and reversible across successive A–B–A reversals (Figure 3B–D).

Representative occupancy heat maps (Figure 3D) illustrate that animals consistently avoided the stimulation-paired compartment during each conditioning phase, with place preference reversing after each reversal of the stimulation contingency. This avoidance was expressed as a context-specific inhibition of entry into the stimulation-paired compartment rather than as overt motor responses or stereotyped defensive behaviors. Statistical analysis confirmed these observations and revealed that the percentage of time spent in the initially preferred compartment differed significantly across experimental phases (baseline, initial conditioning, first reversal, and second reversal; repeated-measures ANOVA: *F*(3,18) = 88,676, *p* < 0.001; Figure 3C). Bonferroni-corrected post hoc comparisons revealed a significant decrease in the time spent in the initially preferred compartment following the initial conditioning phase compared with baseline (p=0.001), a significant increase during the first reversal (p< 0.001), and a subsequent significant decrease during the second reversal (p<0.001). A more detailed analysis of the percentage of time spent in the compartment not associated with the stimulation across 5-min training blocks revealed that animals in the Dmr group showed a progressive increase in time spent in this compartment during both the initial conditioning phase and the first reversal phase. This pattern was supported by the significant effect of training in the repeated-measures ANOVA (first conditioning: F(5,30) = 5.986, p = 0.001; first reversal: F(5,30) = 2.723; p = 0.038) demonstrating effective learning (Figure 3B). An analysis of the dynamics of the transitions between the end of each phase and the beginning of the subsequent phase (5-min blocks), based on the percentage of time spent in the initially preferred compartment, revealed significant differences across all three phase transitions (initial conditioning phase t(6) = 5.336, p < 0.002, the first reversal t(6) = −8.205, p < 0.001, the second reversal t(6) = 8.737, p < 0.001), indicating that the behavioral shifts occurred rapidly (Figure 3B).

Likewise, analysis relative to chance confirmed that Dmr-stimulated animals consistently avoided the stimulation-paired compartment throughout all stimulation phases (first phase: t(6) = −9.848, p < .001; first reversal: t(6) = −10.374, p < 0.001; second reversal: t(6) = −9.176, p < 0.001; Figure 3E), demonstrating that focal activation of Dmr is sufficient to produce robust conditioned place avoidance in freely moving goldfish.

In contrast to Dmr, animals receiving focal microstimulation in Dmc showed no systematic changes in place preference across experimental phases (Figure 3B–D). Place preference remained stable throughout the experiment, and representative occupancy maps revealed no consistent redistribution of spatial exploration following changes in the stimulation contingency (Figure 3D). Consistent with these observations, repeated-measures ANOVA revealed no significant effect of experimental phase (baseline, initial conditioning, first reversal, and second reversal) on the percentage of time spent in the initially preferred compartment (F(3,18) = 0.134, p = 0.939; Figure 3C). Similarly, Dmc-stimulated animals did not differ from chance in the proportion of time spent in the stimulation-paired compartment during any stimulation phase (one-sample *t* tests: first phase, *t*(6) = 1.091, *p* = 0.317; first reversal, *t*(6) = −0.722, *p* = 0.498; second reversal, *t*(6) = 0.583, *p* = 0.581; Figure 3E), indicating that focal activation of Dmc did not generate conditioned place avoidance.

Direct comparison of the two stimulation groups further demonstrated the specificity of this behavioral effect. The magnitude of place preference change between successive phases was significantly greater in the Dmr group than in the Dmc group across all phases (first phase: *t*(12) = 4.321, *p* = 0.001; first reversal: *t*(12) = 6.512, *p* < 0.001; second reversal: *t*(12) = 5.812, *p* < 0.001; Figure 3F). Finally, an avoidance index, calculated by subtracting the time spent in the stimulation-paired compartment from the time spent in the non-stimulated compartment within each stimulation phase and averaged across the three stimulation phases, was also significantly greater in the Dmr group than in the Dmc group (*t*(12) = 6.408, *p* < 0.001; Figure 3G).

Together with the sensory imaging experiments, these findings demonstrate a clear functional dissociation within the rostrocaudal subdivisions of Dm. Sensory-evoked activity is selectively localized to Dmc, whereas focal activation of Dmr is sufficient to generate conditioned place avoidance.

### Anatomical connectivity reveals a distributed circuit linking the Dmc and Dmr pallial domains

The complementary functional roles of Dmc and Dmr revealed by the preceding experiments raise the question of how these two pallial domains are integrated into the broader brain circuits that transform sensory and bodily-related information into motivational control. We therefore asked whether distinct patterns of anatomical connectivity could account for the complementary functional properties of Dmc and Dmr (Figure 4A).

**Figure 4.**
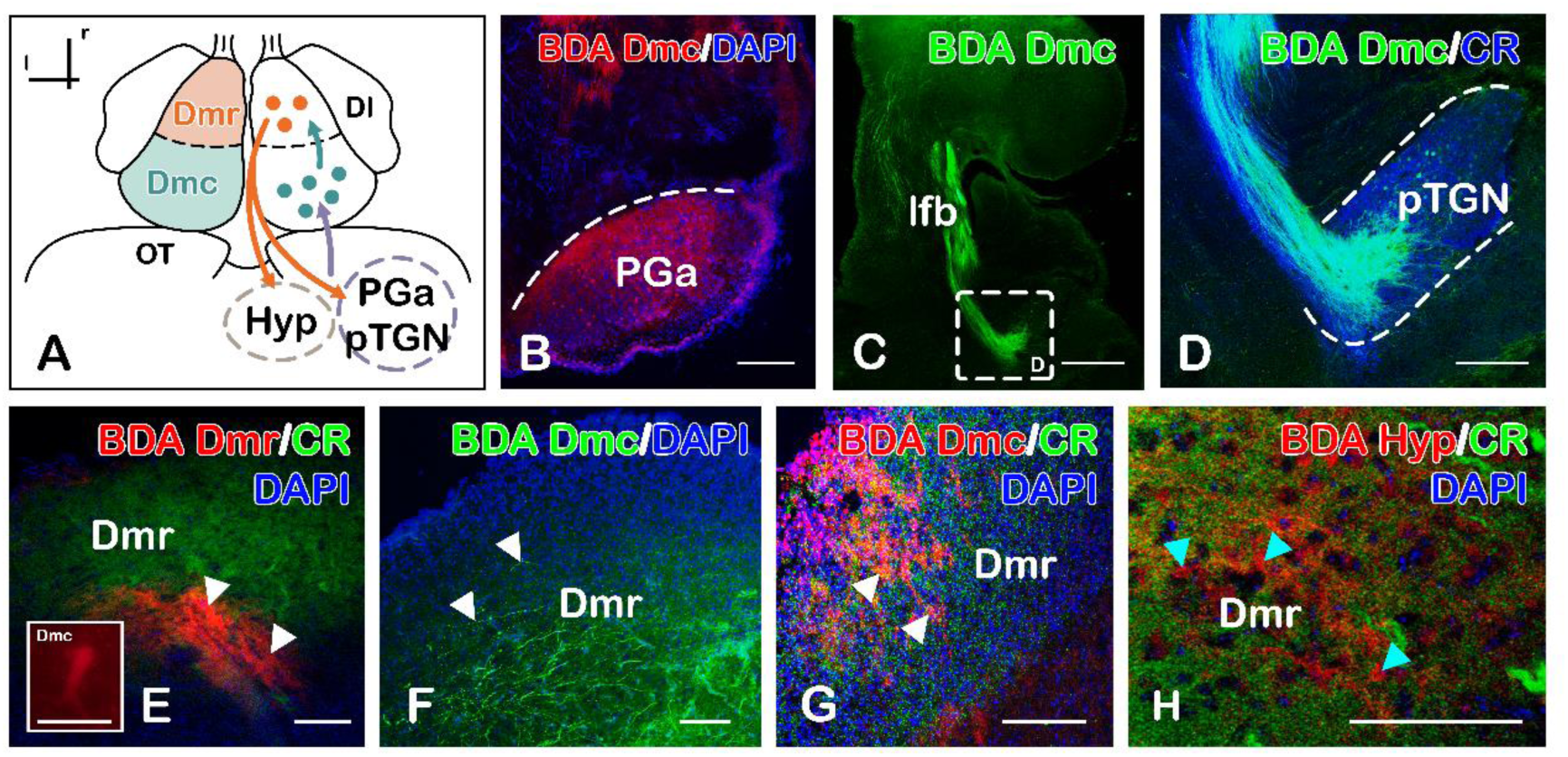
Connectivity of the goldfish dorsomedial pallium reveals a circuit linking sensory inputs to motivational output pathways. **A**. Schematic dorsal view of the goldfish telencephalon summarizing the connectivity between the caudal (Dmc) and rostral (Dmr) subdivisions of the dorsomedial pallium. Color coding indicates the principal inputs and outputs identified in the present study. Large dots represent neuronal cell bodies, and arrows indicate fiber pathways. **B–H**. Representative photomicrographs of transverse brain sections following biotinylated dextran amine (BDA) tracer applications in combination to DAPI and calretinine (CR). The color code for BDA injections and CR is shown in the upper right corner of each panel. *Abbreviations*: Dl, lateral part of the dorsal telencephalic area; Dmc, caudal division of the medial part of the dorsal telencephalic territory; Dmr, rostral division of the medial part of the dorsal telencephalic territory; Hyp, hypothalamus; lfb, lateral forebrain bundle; PG, preglomerular complex; PGa, anterior division of the preglomerular complex; pTGN, preglomerular tertiary gustatory nucleus.

To address this question, we examined the anatomical connectivity of Dmc and Dmr using dextranamine axonal tracing in the goldfish. Tracer injections centered in Dmc, combined with CR immunolabeling to precisely identify the rostrocaudal level of the injected territory, produced strong retrograde labeling in the anterior preglomerular complex (PGa; Figure 4B) and the preglomerular tertiary gustatory nucleus (pTGN), the latter corresponding to CR-positive neurons, together with labeled fibers coursing through the lateral forebrain bundle (Figure 4C,D). Tracer injections centered in Dmr revealed a prominent efferent tract (arrows in Figure 4E) together with retrogradely labeled neurons in Dmc (higher magnification in Figure 4E). Complementary injections into Dmc revealed labeled fibers projecting rostrally toward Dmr (white arrows in Figure 4F,G), demonstrating robust intrapallial connectivity between Dmc and Dmr and confirming a direct projection from caudal Dmc to rostral Dmr. The descending projections arising from Dmr (Figure 4E) and coursing through subpallial territories toward hypothalamic and brainstem targets were further confirmed by complementary tracer injections into the hypothalamic region, combined with CR immunohistochemistry, which produced retrogradely labeled neurons within Dmr (cyan arrowheads in Figure 4H).

Taken together, these anatomical findings support a distributed circuit in which multimodal sensory and bodily-related inputs converge onto Dmc and are relayed to Dmr through direct intrapallial projections. Dmr, in turn, constitutes the principal descending pallial output, projecting to hypothalamic and brainstem centers involved in autonomic and behavioral regulation. This organization provides the anatomical substrate through which sensory-responsive and motivationally specialized pallial domains interact to transform sensory and bodily-related information into adaptive behavioral responses.

**Figure 5.**
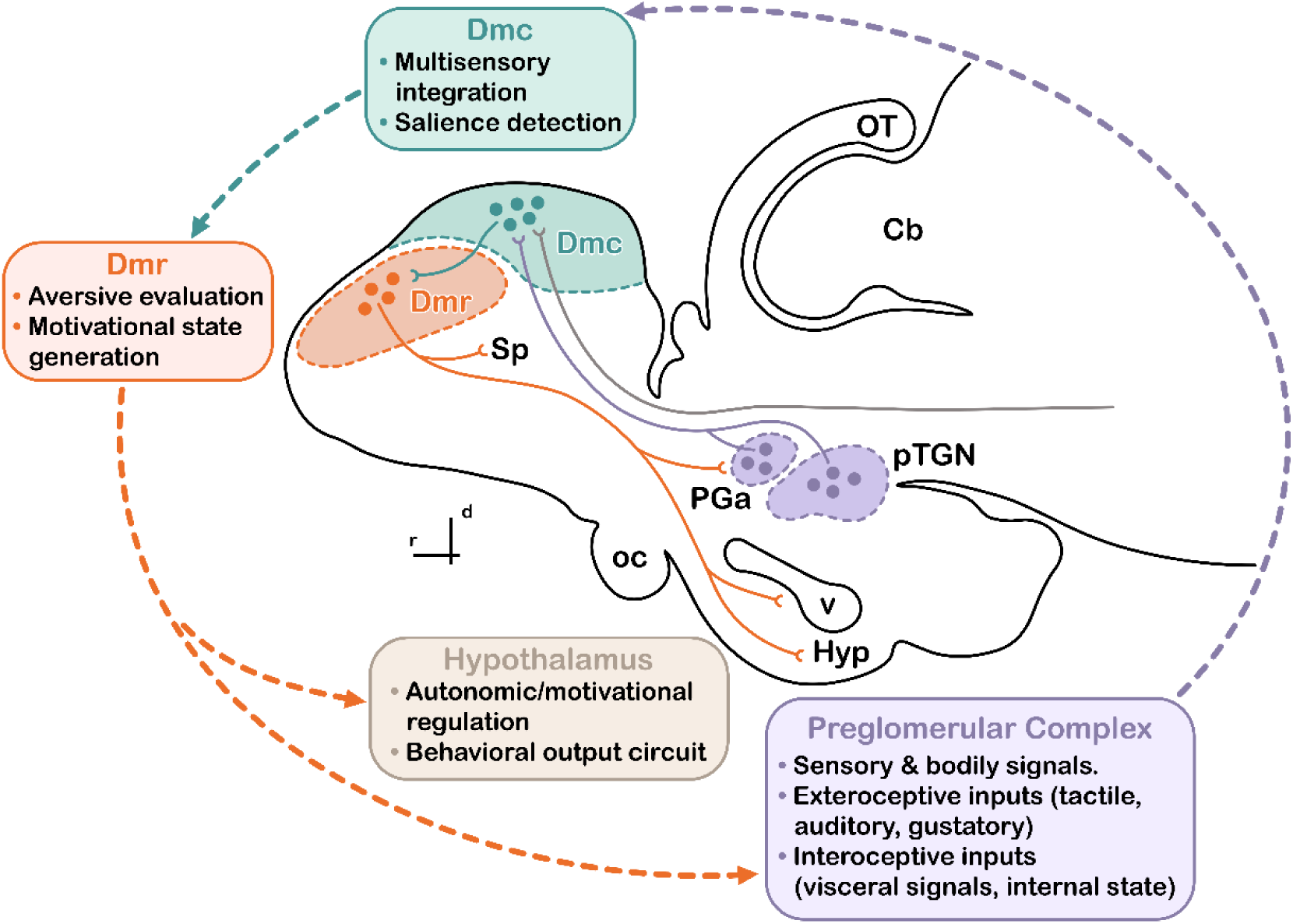
Conceptual diagram of the distributed pallial circuit linking multimodal sensory and bodily representations to aversive motivational control in the goldfish telencephalon. The caudal subdivision of the dorsomedial pallium (Dmc) receives convergent exteroceptive and interoceptive inputs from the preglomerular complex and other diencephalic and brainstem nuclei. Within Dmc, multisensory and bodily signals are integrated and contribute to the evaluation of biologically relevant stimuli. This information is relayed intrapallially to the rostral subdivision (Dmr), where these integrated sensory and bodily-related representations acquire negative motivational value. Dmr provides descending projections to hypothalamic and brainstem circuits involved in autonomic regulation and behavioral control, thereby linking pallial processing to motivationally guided action. Abbreviations: Cb, cerebellum; Dmc, caudal division of the medial part of the dorsal telencephalic area; Dmr, rostral division of the medial part of the dorsal telencephalic area; Hyp, hypothalamus; OT, optic tectum; PG, preglomerular complex; Sp, subpallium.

## DISCUSSION

The present results identify a previously unrecognized distributed functional organization of the teleost dorsomedial pallium (Dm), in which anatomically distinct but interconnected rostrocaudal domains perform complementary computations within a distributed pallial circuit. The caudal domain (Dmc) contains modality-specific sensory representations and receives convergent sensory and bodily-related inputs, consistent with a role in integrating exteroceptive and bodily information into biologically relevant representations. The rostral domain (Dmr), by contrast, exhibits little or no sensory responsiveness under the conditions examined, is sufficient to generate conditioned place avoidance, and constitutes the principal source of descending outputs to hypothalamic and brainstem circuits involved in autonomic and behavioral regulation. These findings indicate that Dm operates as a distributed pallial circuit in which interactions between complementary pallial domains transform integrated sensory and bodily representations into aversive motivational signals capable of guiding flexible instrumental behavior. This functional architecture provides a new framework for understanding both the organization of the teleost dorsomedial pallium and for interpreting its evolutionary relationships with distributed pallial systems in other vertebrates.

### A distributed pallial circuit for sensory–motivational transformation

Our results indicate that the teleost Dm is organized as a distributed pallial circuit composed of two interconnected domains. Previous studies have alternatively emphasized either sensory processing (Prechtl et al., 1998; von der Emde & Prechtl, 1999; Trinh et al., 2026) or emotional control, particularly aversively motivated learning and defensive behavior (Portavella et al., 2004; Broglio et al., 2005; Martín et al., 2011; Lal et al., 2018; Trinh et al., 2026), as defining functions of Dm. Our findings reconcile these apparently contrasting interpretations by showing that these domains perform complementary computations linking sensory and bodily-related integration with motivational valuation and behavioral control.

Wide-field voltage-sensitive dye imaging demonstrates that Dmc contains spatially organized representations of tactile, auditory and gustatory stimuli. Rather than exhibiting diffuse multisensory convergence, sensory responses remain organized into modality-specific territories with selective zones of overlap. Somatosensory activity occupies a central position between gustatory and auditory representations, partially overlapping with both modalities, whereas auditory and gustatory representations remain largely segregated. Together with the recently described somatotopic, tonotopic and gustotopic maps in Dmc (Ocaña et al., 2026), these findings indicate that Dmc preserves modality-specific sensory information while permitting controlled interactions among behaviorally relevant sensory channels, extending previous evidence supporting a sensory integrative role for this region (Prechtl et al., 1998; von der Emde and Prechtl, 1999; Trinh et al., 2026). The central position of somatosensory activity further suggests that body-related information may facilitate interactions among multiple sensory modalities. More generally, these findings are consistent with evidence that somatosensory representations are embedded within distributed multimodal networks supporting object perception, body representation and action (de Haan et al., 2020). They also agree with current theories proposing that adaptive behavior depends on the integration of external sensory information with bodily state and interoceptive signals, thereby providing the physiological context for perception, cognition and action (Damasio, 1999; Craig, 2009; Barrett & Simmons, 2015; Pessoa, 2017; Quadt et al., 2018).

The connectivity of Dmc provides additional support for this interpretation. Dmc receives dense multisensory afferents from the preglomerular complex, the principal ascending sensory relay to the teleost pallium, conveying auditory, gustatory and somatosensory information through distinct preglomerular subnuclei (Northcutt, 2006; Wullimann and Mueller, 2004; Yamamoto and Ito, 2005; Yamamoto et al., 2007; Mueller et al., 2012; Kato et al., 2012; Striedter, 2021; Yáñez et al., 2021; Trinh et al., 2026). In addition, Dmc receives inputs from diencephalic and brainstem regions associated with visceral, homeostatic and bodily signaling (Giassi et al., 2012; Bloch et al., 2020; Gutjahr et al., 2025), whereas the preglomerular complex itself receives projections from hypothalamic and preoptic nuclei, providing an additional indirect route through which internal-state information can reach Dmc (Giassi et al., 2012). These anatomical and physiological observations indicate that Dmc is strategically positioned to integrate exteroceptive and interoceptive information into neural representations reflecting the current biological significance of environmental events. Dmc may construct representations of the external environment within the context of the animal’s current physiological condition, providing the informational substrate upon which Dmr may compute aversive motivational value.

In striking contrast, Dmr exhibited little or no detectable sensory-evoked activity under the present stimulation conditions. Focal electrical microstimulation consistently induced robust, reversible and flexible conditioned place avoidance without producing stereotyped defensive behaviors, whereas equivalent stimulation of Dmc failed to generate comparable behavioral effects. Together with its prominent descending projections to subpallial, hypothalamic, preoptic and brainstem regions (Folgueira et al., 2004; Northcutt, 2006; Giassi et al., 2012; Lal et al., 2018; Yáñez et al., 2021), these findings identify Dmr as the principal source of aversive motivational signals within the Dm network. Rather than directly encoding sensory information, Dmr appears to transform biologically relevant representations into aversive value, thereby influencing behavior through the recruitment of conserved autonomic and behavioral effector systems. The aversive motivational signals generated by Dmr may also influence other pallial networks, including the dorsolateral pallium (Dl), a hippocampal-like pallial region supporting spatial, relational and episodic-like memory, and thus contribute to the formation and updating of adaptive memory representations (Rodríguez et al., 2002; Broglio et al., 2005; Salas et al., 2006; McGaugh, 2004; Pessoa, 2017). Previous lesion studies demonstrated that damage to Dm disrupts avoidance learning, conditioned autonomic responses and defensive behaviors (Portavella et al., 2004; Broglio et al., 2005; Martín et al., 2011), but because these experiments targeted Dm as a whole, they could not determine they could not determine whether the entire Dm contributed uniformly to these functions or whether they were preferentially mediated by specific subregions. The present findings refine this interpretation by showing that these motivational functions emerge from complementary interactions between Dmc and Dmr, while identifying Dmr as the pallial domain selectively involved in assigning aversive motivational value.

The characteristics of the place aversion conditioning procedure used in this study further clarify the functional significance of this motivational system. Unlike conventional conditioned place aversion procedures, in which aversive outcomes are paired with contextual cues independently of the animal’s ongoing behavior, intracerebral stimulation in the present task was contingent upon the fish’s own spatial choices. Entry into the designated compartment initiated stimulation, whereas leaving it immediately terminated it, establishing an instrumental action–outcome contingency. Within classical theories of instrumental learning, outcomes acquire behavioral significance by modifying the future probability of actions through their reinforcing or punishing consequences (Thorndike, 1911; Dickinson, 1985). Early in each training phase, termination of Dmr stimulation reinforced escape from the stimulated compartment; with continued experience, fish progressively prevented further stimulation by withholding entry into that compartment and remaining in the alternative one. The resulting behavior therefore consisted of an instrumental form of passive avoidance, indicating that Dmr stimulation effectively guided the acquisition of instrumental avoidance behavior (Rolls, 1999, 2014). The repeated reversals further demonstrate that the behavioral consequences of Dmr activation are not tied to a particular defensive response. At the beginning of each reversal, fish initially escaped from the newly stimulated compartment but, with continued training, progressively returned to passive avoidance. This passive avoidance strategy was reacquired increasingly rapidly across successive reversals, indicating that Dmr activation flexibly supports different behavioral responses according to the prevailing action–outcome contingency rather than specifying a fixed defensive strategy.

The present findings also help reinterpret the pioneering studies that first identified the teleost telencephalon as essential for instrumental avoidance learning. Early lesion experiments demonstrated that complete telencephalic ablation selectively impaired passive avoidance (Overmier & Flood, 1969) and disrupted performance during successive reversal learning (Frank, Flood & Overmier, 1972), while leaving classical conditioning and many other forms of learning relatively preserved (Hainsworth et al., 1967; Kaplan & Aronson, 1969; Overmier & Curnow, 1969; Savage, 1969; Flood & Overmier, 1971). On the basis of these findings it has been proposed that the teleost telencephalon functioned as a limbic-like forebrain system specialized for instrumental motivated behavior (Flood, Overmier & Savage, 1976). Subsequent lesion studies localized these functions to the dorsomedial pallium (Rodríguez et al., 2002; Portavella et al., 2004; Broglio et al., 2005; Martín et al., 2011). The present findings extend this functional framework by identifying complementary roles for Dmc and Dmr within the distributed pallial circuit mediating instrumental avoidance learning.

### Comparative and evolutionary considerations: A distributed pallial circuit rather than a pallial amygdala

The present findings have important implications for understanding the evolution of vertebrate pallial organization. They reconcile the apparently disparate sensory and aversive functions attributed to the teleost dorsomedial pallium (Dm) and provide a new framework for interpreting its evolutionary significance. Rather than supporting a simple one-to-one correspondence between the teleost Dm and a single mammalian pallial structure, our results indicate that Dm is organized as a distributed functional network in which anatomically distinct but strongly interconnected rostrocaudal domains perform complementary computational operations, separating sensory integration from motivational valuation while maintaining strong anatomical and functional coupling between the two domains. Comparable principles of functional organization characterize mammalian mesocortical limbic networks, in which insular, cingulate and ventromedial prefrontal cortices interact to integrate interoceptive and exteroceptive information, assign motivational value, and guide adaptive behavior through distributed corticolimbic networks rather than within a single anatomical structure (Mogenson et al., 1980; Mesulam, 1998; Swanson, 2000; Craig, 2009; Puelles et al., 2024; Pessoa, 2013, 2017; Cisek, 2019, 2022).

This framework provides a new basis for evaluating competing hypotheses concerning the homologies of Dm. Some models have proposed that Dm constitutes the pallial component of the teleost amygdaloid complex (Lal and Kawakami, 2022), whereas Porter and Mueller (2020) suggested more specifically that caudal Dm corresponds to a posteromedial pallial amygdaloid area. In addition, developmental studies have firmly established the existence of a subpallial extended amygdala in teleosts (Biechl et al., 2017; Mueller, 2022; Armbruster et al., 2025). Using the conserved developmental markers Otp, Pax6 and tyrosine hydroxylase, we identified the neighboring subpallial amygdaloid territories, whereas both Dmc and Dmr remained molecularly distinct from these territories and lacked the combination of developmental features expected for an amygdaloid region. Thus, our results support the existence and anatomical extent of the teleost subpallial extended amygdala, but do not support the hypothesis that either rostrocaudal subdivision of Dm constitutes its pallial counterpart.

This framework provides a new basis for evaluating competing hypotheses concerning the homologies of Dm. Developmental studies have firmly established the existence of a subpallial extended amygdala in teleosts (Biechl et al., 2017; Mueller, 2022; Armbruster et al., 2025). Broader Developmental studies have firmly established the existence of a subpallial extended amygdala in teleosts (Biechl et al., 2017; Mueller, 2022; Armbruster et al., 2025). Some models have proposed that Dm constitutes the pallial component of the teleost amygdaloid complex (Lal and Kawakami, 2022), whereas Porter and Mueller (2020) suggested more specifically that caudal Dm corresponds to a posteromedial pallial amygdaloid area. In addition, dDevelopmental studies have firmly established the existence of a subpallial extended amygdala in teleosts (Biechl et al., 2017; Mueller, 2022; Armbruster et al., 2025). Using the conserved developmental markers Otp, Pax6 and tyrosine hydroxylase, we identified the neighboring subpallial amygdaloid territories, whereas both Dmc and Dmr remained molecularly distinct from these territories and lacked the combination of developmental features expected for an amygdaloid region. Thus, our results support the existence and anatomical extent of the teleost subpallial extended amygdala, but do not support the hypothesis that either rostrocaudal subdivision of Dm constitutes its pallial counterpart.

This interpretation is further supported by recent transcriptomic evidence. Single-cell and spatial transcriptomic analyses of the goldfish telencephalon (Tibi et al., 2023) revealed pronounced molecular heterogeneity within Dm, identifying distinct glutamatergic neuronal populations distributed along its rostrocaudal axis. Specifically, anterior Dm is enriched in neuronal populations expressing genes such as NEUROD1 and CNR1, whereas more posterior regions contain alternative glutamatergic populations associated with NR2F2. This molecular organization closely parallels the functional subdivision identified here. However, despite revealing extensive cellular diversity, these analyses do not identify a discrete neuronal population corresponding to a pallial amygdaloid nucleus within Dm. Likewise, the rostrocaudal differences in calretinin immunoreactivity observed here, consistent with previous reports (Castro et al., 2006), together with additional markers defining the pallial-subpallial boundary (Figure 1; Tibi et al., 2023), support the interpretation that Dm is internally regionalized into functionally differentiated pallial domains rather than containing a discrete pallial amygdaloid territory.

The functional differentiation of Dm revealed here is therefore more consistent with a distributed paralimbic architecture than with the functional organization expected from a unitary pallial amygdala or a neocortical sensory hierarchy. Rather than simply exhibiting isolated functional similarities to individual mammalian structures, the Dmc–Dmr organization reproduces a broader computational arrangement in which the construction of biologically meaningful sensory–bodily representations is segregated from the assignment of motivational value while both processes remain tightly integrated within a common functional circuit. Within this framework, Dmc occupies a position appropriate for integrating exteroceptive and bodily information into biologically meaningful representations, closely resembling the integrative role of the mammalian insular cortex within distributed corticolimbic networks. Dmr, in contrast, transforms these representations into behavior-contingent aversive value, supports reinforcement-guided updating of instrumental behavior, and recruits descending autonomic and behavioral control systems, functions more commonly associated with anterior cingulate, ventromedial prefrontal, striatal and related corticolimbic circuits (Rolls, 2014; Pessoa, 2017).

More broadly, these results suggest that one of the fundamental organizational principles underlying affective processing may have emerged early during vertebrate evolution. Rather than relying on a single multifunctional pallial center, adaptive behavior appears to depend on interactions among functionally specialized but anatomically interconnected pallial territories, in which the construction of biologically meaningful sensory–bodily representations is separated from the assignment of aversive motivational value while preserving their close functional integration. Rather than merely representing a simplified version of the mammalian forebrain, the teleost Dm may therefore constitute an early vertebrate implementation of a distributed computational architecture that could have been conserved across vertebrate evolution.

## MATERIAL AND METHODS

### Animals

Adult goldfish (Carassius auratus) were used in all experiments. For optical imaging experiments, six animals measuring 12–15 cm in length were used. Fourteen animals, measuring 14–16 cm in length, were used for intracerebral microstimulation and behavioral experiments, and eight were used for neuroanatomical and connectivity analyses. Fish were housed in aerated freshwater tanks under controlled conditions (20°C; 12:12 h light–dark cycle) and fed twice daily with dry food pellets (TetraPond PondSticks, Germany). Animals were acclimatized to laboratory conditions for at least one week prior to experimental procedures. All procedures were conducted in accordance with Directive 2010/63/UE of the European Community Council and Spanish legislation (R.D. 53/2013).

### Optical imaging of evoked sensory activity

#### Animal preparation

Goldfish were anesthetized by immersion in a 1:20,000 solution of tricaine methanesulfonate (MS-222; Sigma-Aldrich), adjusted to pH 7.0–7.5, and anesthesia was maintained by continuous perfusion of aerated anesthetic solution through the gills. The dorsal skin and skull over the telencephalon were removed under microscopic guidance. Intracranial fatty tissue and the tela choroidea were carefully removed to expose the dorsal telencephalic surface. After surgery, the anesthetic solution was replaced with fresh water until spontaneous respiration and normal movements resumed.

#### Voltage-sensitive dye staining

The voltage-sensitive dye Di-2-ANEPEQ (JPW 1114; Molecular Probes) was used. A working solution (50 μg/ml) was prepared by diluting a stock solution (0.5 mg/ml in distilled water) in goldfish Ringer solution (116 mM NaCl, 2.9 mM KCl, 1.8 mM CaCl₂, 5 mM HEPES, pH 7.2). Approximately 100 μl of dye solution was applied to the exposed telencephalon for 45 minutes. The tissue was then rinsed twice with Ringer solution to remove excess dye and maintain hydration. During recording sessions, animals were immobilized by intramuscular injection of d-tubocurarine (0.002 mg/g; Merck) to eliminate movement artifacts.

#### Sensory stimulation

Three sensory modalities were tested. Tactile stimulation consisted of a brief water jet (0.5 ml/s, 200 ms) applied to the left side of the trunk at the rostral region of the dorsal fin insertion. The jet was delivered using a programmable syringe pump (Aladdin; World Precision Instruments) through a needle positioned 0.3–0.5 cm from the skin surface. Auditory stimulation consisted of pure tones (200 ms duration, 0.5 kHz, 90 dB, abrupt rise and fall times) generated by an auditory stimulator (LE-150; Letica Scientific Instruments) and delivered through a loudspeaker positioned approximately 50 cm behind and above the animal’s head. Gustatory stimulation was delivered via a programmable syringe pump (Aladdin; World Precision Instruments) at a flow rate of 0.5 ml/s for 1 s into the water flow bathing the oral cavity. Silicone tubes (4 mm diameter) converged into a main tube inserted into the animal’s mouth. The tastant was a 2-phenylethanol solution (10⁻² mol/L in distilled water). Water in the experimental chamber was continuously replaced to prevent chemical accumulation.

#### Optical recording and analysis

Optical recordings were obtained using a MiCAM01 imaging system (Brain Vision) coupled to an epifluorescence microscope (THT; Scimedia) mounted on a vibration-isolation table. Excitation was provided by a 150 W tungsten–halogen lamp (MHF-G150LR; Moritex) through a 530 ± 3 nm excitation filter and a dichroic mirror. Emitted fluorescence (>590 nm) was captured using a CCD camera (90 × 60 pixels), covering a 2.9 × 2.1 mm imaging area. With a 0.63× objective (PLAN APO; Leica Microsystems) and a 1× projection lens, the effective field of view was 4.6 × 3.3 mm, centered on the dorsal telencephalon.

Images were acquired at 6 ms per frame (166.7Hz). Each trial lasted 3,400 ms, including a 300 ms prestimulus baseline period. Recordings began 300 ms after illumination onset to allow stabilization of fluorescence signals. Fifteen trials were collected for each sensory modality with intertrial intervals of 30-60 s and averaged to improve signal-to-noise ratio. Image processing was performed using BV-Analyzer software (Brain Vision). Changes in optical intensity (dF) were quantified as the difference of fluorescence in each moment (F) in relation to the fluorescence of the reference images (F0), calculated as the mean fluorescence in the 30 photograms before stimulus onset. Signals were detrended to correct for bleaching, spatially filtered using a 7 × 7 pixel Gaussian filter, and temporally smoothed using a low-pass filter. Pseudocolor activity maps displayed dF and were thresholded at 25% of the full-scale range. Upward deflections in time traces represent depolarization. Data were normalized by the maximal signal amplitude.

### Chronic intracerebral microstimulation and conditioned place aversion

#### Surgery and electrode implantation

Animals were anesthetized with MS-222 (1:20,000, buffered to tank pH) and placed in a custom surgical tank. A continuous flow of aerated anesthetic solution was delivered to the gills via an oral cannula. The skin over the skull was incised and disinfected, the periosteum removed, and a craniotomy performed over the telencephalon. A stainless-steel screw serving as indifferent electrode was inserted at the anterior margin of the craniotomy.

Fat covering the telencephalon was removed by gentle aspiration, and the meninges were retracted. Stainless-steel microelectrodes (25 μm diameter, glass-insulated, impedance 100–700 kΩ) were inserted under visual control using a micromanipulator (Narishige) to depths of 70–250 μm, targeting either the rostral (Dmr) or caudal (Dmc) medial pallium. Electrodes were secured with dental cement, and the connectors were insulated with silicone.

Electrode locations were recorded during surgery by registering the anteroposterior, mediolateral, and dorsoventral coordinates of each implantation site on a standard atlas of the goldfish telencephalon. In addition, photographs of the implanted electrodes were taken in all animals to document their position relative to external brain landmarks. In some subjects, electrode locations were further verified histologically by producing a small electrolytic lesion (10 μA DC for 10 s). Animals were euthanized by an overdose of MS-222 and transcardially perfused with PBS followed by a methanol–acetone–water fixative. Brains were embedded in paraffin, sectioned transversely, stained with cresyl violet (Nissl), and examined to confirm electrode locations.

#### Behavioral apparatus

Behavioral experiments were conducted in a glass tank (83 × 83 × 27 cm) lined with white PVC and placed in a sound- and light-isolated room. Two colored plastic sheets (light blue and pink; 63 × 41.5 cm) divided the tank floor into two compartments. Behavior was recorded using a ceiling-mounted webcam connected to a PC running AnyMaze tracking software (Stoelting). The tracking system automatically triggered stimulation via an AMi-2 interface linked to a stimulator (Cygnus P400) and constant-current isolation unit (SIU9A; A-M Systems). Stimulation parameters were monitored on a digital oscilloscope (Tektronix).

#### Microstimulation protocol

Electrical stimulation consisted of 2-s trains of 50 Hz pulses, with each pulse lasting 200 μs. Stimulation intensity was set individually for each electrode at 25% below the motor response threshold. For Dmr electrodes, intensities ranged from 160–220 μA; for Dmc electrodes, from 180–240 μA.

The stimulation intensity for each electrode was individually adjusted by gradually increasing the current until a clearly observable behavioral response was elicited without inducing escape behavior. The current was then reduced by 25%, and this value was used as the experimental stimulation intensity.

#### Conditioned place aversion procedure

The experiment consisted of a baseline phase and three stimulation phases of 30-min separated by 5-min rest periods. Animals then underwent a conditioned place avoidance paradigm using a serial A–B–A reversal design, in which the stimulation-paired compartment was alternated across successive phases. For each fish, the compartment selected for stimulation during the first conditioning phase was its preferred compartment during the baseline period, defined as the compartment in which the fish spent more than 50% of the baseline period. Stimulation was automatically applied each time the animal’s head entered the designated compartment and every 5 s while it remained inside. The experimenter was blinded to the region in which the electrode was located.

Time spent in each compartment was quantified for each phase. The magnitude of place preference change between successive phases was calculated by subtracting the percentage of time spent in the stimulation-paired compartment during the current phase from the percentage of time spent in the same compartment during the immediately preceding phase, when it was not paired with stimulation. Positive values indicate a shift in preference toward the newly non-stimulation-paired compartment, whereas values close to zero indicate little or no change. An avoidance index was calculated by subtracting the time spent in the stimulation-paired compartment from the time spent in the non-stimulated compartment within each stimulation phase. Positive values indicate avoidance of the stimulation-paired compartment, whereas values close to zero indicate no preference between compartments.

#### Statistical analysis

Statistical analyses were performed using SPSS (IBM). Data are reported as mean ± SEM. For optical imaging experiments, comparisons between Dmc and Dmr responses were performed using paired-sample *t* tests, whereas comparisons of responses across the different sensory modalities were performed using repeated-measures ANOVA, followed by Bonferroni-corrected post hoc comparisons when appropriate. Behavioral measures in the conditioned place aversion experiment were analyzed using repeated-measures ANOVA to assess changes across experimental phases, followed by Bonferroni-corrected post hoc comparisons when appropriate. Planned paired-sample t tests were additionally performed to compare specific experimental phases, whereas independent-sample t tests were used for comparisons between independent groups, as appropriate. Preference values were additionally compared against chance level (50%) using one-sample t tests. Statistical significance was set at p < 0.05.

### Neuroanatomical analysis

#### Immunofluorescence

After being deeply anesthetized (0,1% MS222, pH 7.4; Sigma-Aldrich Merck KGaA, Darmstadt, Germany), the animals were perfused transcardially with 0.9% NaCl saline solution, followed by cold 4% paraformaldehyde, (in a 0.1 M phosphate buffer; pH 7.4) fixative solution. The brains were cryoprotected in a solution of 30% sucrose in PB (0.1M phosphate buffer pH 7.4) for 4-6 hours at 4°C and cut on freezing microtome in 40 µm transverse and sagittal slices.

Single and double immunolabeling experiments were conducted on free-floating sections, using the primary antibodies: mouse anti-CR (1:500; Swant; Cat. No. 6B3); rabbit anti-Ctip2 (1:100; Absolute Antibody; Cat. No. Ab00616-23.0); rabbit anti-GABA (1:500; Merck; Cat. No. A2052); rabbit anti-GFAP (1:500; DAKO-Agilent; Cat. No. Z0334); mouse anti-Meis2 (1:50; DSHB; Cat. No. PCRP-MEIS2-1A11); rabbit anti-Otp (1:200; PickCell); rabbit anti-Pax6 (1:300; Abcam; Cat. No. AB5790-100) and mouse anti-TH (1:1.000; Immunostar; Cat. No. 22941). They were diluted in PB containing 0.5% Triton X-100, as follows: the sections were incubated with primary antibody for 48 h at 4°C, and after being rinsed in PB they were incubated with the secondary antibody, Alexa 488-conjugated goat anti-mouse antibody (1:300; Thermo Fisher Scientific) and/or Alexa 594-conjugated goat anti-rabbit antibody (1:300; Thermo Fisher Scientific) for 2 h at room temperature.

Antibodies controls included omission and incubation with preimmune mouse or rabbit sera instead of the primary antibody. No residual staining was observed in any case. After the immunohistochemical steps, the free-floating sections were mounted. All slides were then coverslipped with a fluorescence mounting medium containing 1.5-µg/ml 4’,6-diamidino-2-phenylindole for DNA counterstaining (UltraCruz, SC-24941, Santa Cruz, Dallas, TX, USA).

#### Axonal tract tracing

For axonal tracing experiments, once deeply anesthetized (0,1% MS222, pH 7.4; Sigma-Aldrich Merck KGaA, Darmstadt, Germany), animals were transcardially perfused with iced oxygenated Ringer’s solution (128.10 mM NaCl, 2.55 mM KCl, 3.4 mM CaCl2·2H2O, 0.2 mM NaHCO3, 10 mM glucose, Merck) equilibrated with carbogen gas (95% O₂, 5% CO₂) to a final pH of 7.4. After perfusion, the brains were dissected, and the dura mater and choroid plexus were removed. Biotinylated dextran amine (BDA 3.000; Thermo Fisher Scientific; Cat. No. D7135) was applied at specific brain regions. Brains were then incubated for 24 hours at 4 °C in continuously oxygenated Ringer’s solution. Following incubation, brains were fixed in 4% paraformaldehyde in 0.1 M phosphate buffer 24 hours. After fixation, tissue was cryoprotected, sectioned as previously described, and processed for tracer visualization. BDA was visualized by incubation with one of the following streptavidin-conjugated fluorophores for 90 minutes: Oregon Green-streptavidin complex (1:500; Thermo Fisher Scientific; Cat. No. S6368) or Alexa 594-streptavidin complex (1:500; Thermo Fisher Scientific; Cat. No. S11227). In all cases, BDA was identified in combination with CR (see above) to precisely determine the location of the injections site and the retrograde/anterograde labeling obtained.

#### Imaging

The sections were analyzed with the microscopes Olympus BX51 microscope, Olympus FV 1200 and Leica sp-2 AOBS confocal. The figure preparation was done with Fiji-Image J (Schindelin et al., 2012), Adobe Photoshop CS6 (Adobe Systems, San Jose, CA) and Canvas X (ACD Systems, Canada).

## Acknowledgements

The authors wish to express their sincere gratitude to María Salud Ramos Guelfo, Mónica Cadena and Eduardo Cueto for their outstanding technical assistance.

## Author contributions

Conceptualization, study design, methodology and funding acquisition: C.S., F.R., N.M.

Voltage-sensitive dye imaging of pallial sensory-evoked responses: C.S.P., B.Q., A.G., A.C., F.R., C.S.

Conditioned place aversion by electrical microstimulation: C.S.P., B.Q., A.G., F.R., C.S.

Immunohistochemistry and tract-tracing: A.C., C.S.P., D.L., J.M.L, N.M. Writing - Original draft: C.S., N.M., F.R.

Writing - Review and editing: C.S., N.M., F.R., C.S.P., B.Q., A.G., A.C., D.L., J.M.L.

## Data availability

Data will be made available on request.

